# A Facile LC-MS Method for Profiling Cholesterol and Cholesteryl Esters in Mammalian Cells and Tissues

**DOI:** 10.1101/2024.04.02.587668

**Authors:** Aakash Chandramouli, Siddhesh S. Kamat

## Abstract

Cholesterol is central to mammalian lipid metabolism and serves many critical functions in the regulation of diverse physiological processes. Dysregulation in cholesterol metabolism is causally linked to numerous human diseases, and therefore, *in vivo*, the concentrations and flux of cholesterol and cholesteryl esters (fatty acid esters of cholesterol) are tightly regulated. While mass spectrometry has been an analytical method of choice for detecting cholesterol and cholesteryl esters in biological samples, the hydrophobicity, chemically inert nature and poor ionization of these neutral lipids has often proved a challenge in developing lipidomics compatible liquid chromatography-mass spectrometry (LC-MS) methods to study them. To overcome this problem, here, we report a reverse-phase LC-MS method that is compatible with existing high-throughput lipidomics strategies, and capable of identifying and quantifying cholesterol and cholesteryl esters from mammalian cells and tissues. Using this sensitive yet robust LC-MS method, we profiled different mammalian cell lines and tissues, and provide a comprehensive picture of cholesterol and cholesteryl esters content in them. Specifically, amongst cholesteryl esters, we find that mammalian cells and tissues largely possess monounsaturated and polyunsaturated variants. Taken together, our lipidomics compatible LC-MS method to study this lipid class opens new avenues in understanding systemic and tissue-level cholesterol metabolism under various physiological conditions.

TOC Graphic

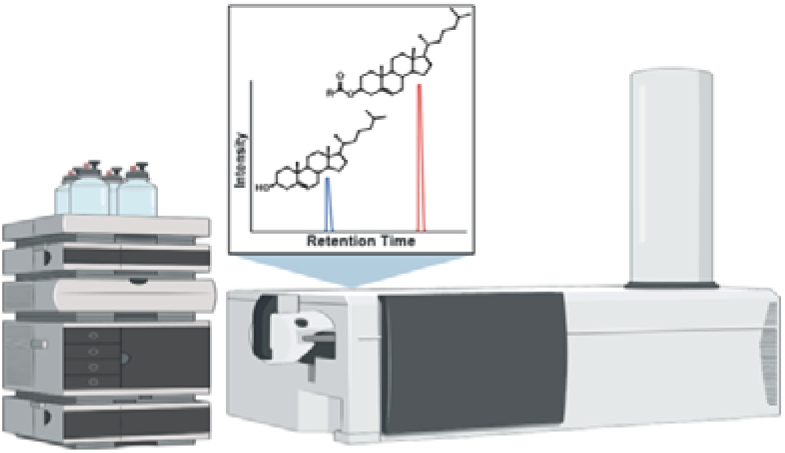

## INTRODUCTION

Cholesterol is a ubiquitous waxy substance (lipid) known to be present in all tissues of our body, and serves many important functions in diverse physiological processes^1–3^. In mammals, cholesterol is systemically obtained via its *de novo* biosynthesis in the liver^4, 5^ or from its exogenous uptake through diet^6–8^. Given its shape and hydrophobicity, cholesterol is an integral part of all cellular membranes in higher organisms (including humans), and regulates the fluidity of these membranes by forming local microdomains termed “lipid rafts”^9, 10^. Cholesterol-rich lipid rafts influence processes such as membrane protein sorting and trafficking, cell polarization, cellular signal transduction pathways and the intracellular transportation of cargo, and have been extensively studied in the context of diverse immunological responses in humans^11, 12^. In addition to the aforementioned processes, cholesterol also serves as a precursor for the biosynthesis of some important biomolecules such as bile salts, steroid hormones, vitamin D to list a few^1, 2, 13^.

*De novo* cholesterol biosynthesis is an energetically expensive process, and hence, mammals have evolved mechanisms to store surplus “free” cholesterol as fatty acid esters of cholesterol (also known as cholesteryl esters). Specifically, two enzymes belonging to the membrane-bound O-acyltransferase (MBOAT) family,^14, 15^ namely acyl-CoA:cholesterol acyltransferase (ACAT) (also known as sterol O-acyltransferase, SOAT)^16, 17^ and lecithin:cholesterol acyltransferase (LCAT),^18–20^ catalyse the conversion of cholesterol to the chemically inert cholesteryl esters in different mammalian tissues (**Figure 1**). ACATs have two isoenzymes: ACAT1 having ubiquitous expression in all mammalian tissues, and ACAT2 having restricted expression in the liver and intestines. Physiologically, both ACATs are responsible for converting excess tissue resident cholesterol to cholesteryl esters, and facilitating their eventual storage as lipid droplets in different mammalian tissues^16, 17^. On the other hand, in mammals, LCAT is very abundant in the blood (or plasma), where it converts excess circulating cholesterol to cholesteryl esters, promotes the formation of high-density lipoproteins (HDLs) via its enzymatic activity and aids reverse cholesterol transport^18–21^. While significant information is available on the formation and storage of cholesteryl esters, little remains known on the identity of the lipases (cholesterylesterases) that metabolize cholesteryl esters back to cholesterol in mammals (**Figure 1**), despite the mention of such an enzymatic activity in literature several years ago^22, 23^.

Nonetheless, due to their involvement in numerous biological processes, the physiological concentrations of cholesterol and cholesteryl esters are tightly regulated^1, 2, 4, 24^, and dysregulation in this metabolism is causally associated with several pathophysiological conditions and diseases in humans^25–34^.

**Figure 1.**
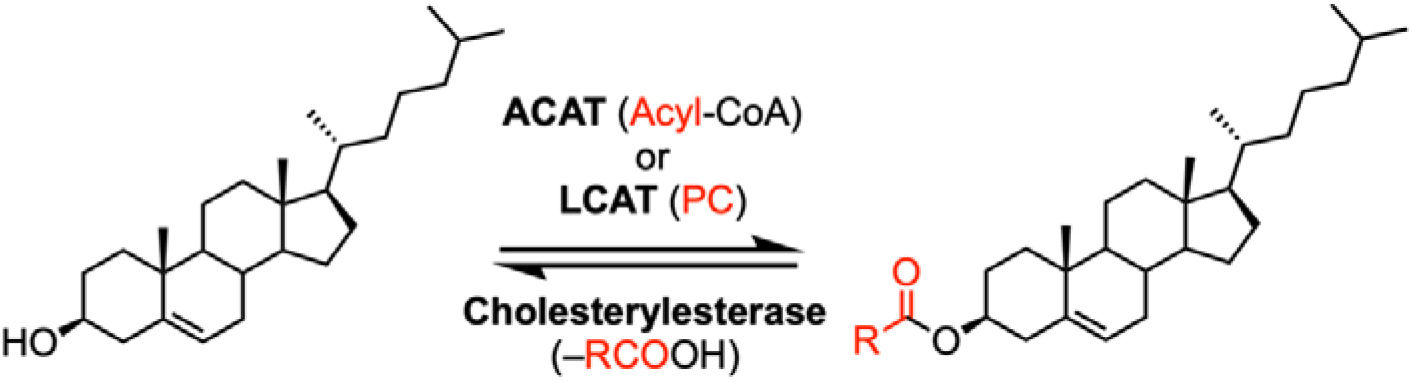
Enzymatic pathways for the metabolism of cholesteryl esters. ACAT and LCAT biosynthesize cholesteryl esters from free cholesterol. Highlighted in red in the parenthesis for ACAT or LCAT is the source of the fatty acid from the second lipid substrate (acyl-CoA for ACAT, and phosphatidylcholine (PC) for LCAT) for these enzymes for generating cholesteryl esters. On the other hand, a cholesterylesterase generates free cholesterol from cholesteryl esters. Highlighted in red in parenthesis for cholesterylesterase is the fate of the acyl-group that eventually forms a free fatty acid, upon the hydrolytic reaction by these enzymes.

Given the importance of cholesterol and cholesteryl esters in human pathophysiology and disease, knowing their physiological concentrations has important clinical significance. Hence, over the years, significant efforts have been put in, towards developing analytical methods and tests to quantify cholesterol and its fatty acid esters in biological systems^35^. Amongst such methods, the first were, a couple of colorimetric tests used to detect total cholesterol content in biological samples such as the blood or serum. In the modified Abell-Kendall method^36^, cholesteryl esters are converted to “free” cholesterol via a saponification reaction using hydroxide, and subsequently the cholesterol in the biological sample is chemically converted to colored compound cholestahexane sulfonic acid (λ_max_ ∼ 410 nm) by acetic-anhydride and sulfuric acid. On similar lines, a few enzyme coupled fluorometric assays have also been developed^37–39^, where cholesteryl esters are first enzymatically converted to “free” cholesterol by using promiscuous esterases (lipases), and subsequently cholesterol in the biological sample is converted to cholest-4-ene-3-one by the action of the enzyme cholesterol oxidase. A byproduct of the cholesterol oxidase catalyzed reaction is hydrogen peroxide^40, 41^, which is then coupled to various sensitive and stable fluorescent probes by enzymatic activities of diverse peroxidases to yield fluorescence signals that can be used to quantify total cholesterol content in biological samples. While rapid and quantitative, a major drawback of both these colorimetric methods is that they provide no knowledge qualitatively or quantitatively of the cholesteryl ester content in biological samples. To circumvent this issue, over the years, a few elegant gas chromatography coupled to mass spectrometry (GC-MS) protocols have been reported^42–47^, that can be used for the determination of free or esterified or total cholesterol (depending on the nature of the pre-treatment step) in biological samples. Despite its quantitative abilities, the major limitations for GC-MS are the cumbersome sample preparation steps due to chemical derivatization of cholesterol and its fatty acid esters prior to analysis, extensive sample run times, poor sensitivity, and extensive need for labor resource. In addition to GC-MS, there are also a few matrix-assisted laser desorption and ionization-based mass spectrometry (MALDI-MS) detection and imaging techniques reported for cholesterol and lipoprotein associated lipids in various biological samples, including tissues^48–52^. While imaging MALDI-MS provides spatial resolution for cholesterol (not well developed for cholesteryl esters yet) in mammalian tissues, this analytical technique has the same limitations as GC-MS.

Lately, the advent of liquid chromatography coupled to mass spectrometry (LC-MS) leveraging electrospray ionization (ESI) techniques has resulted in tremendous progress towards developing high-throughput lipidomics methods with improved speed and sensitivity for detecting and quantifying diverse cellular lipids^53–57^. Yet, even by advanced LC-MS analysis, the measurements of cholesterol and its fatty acid esters from biological samples have remained challenging due to their poor ionization ability, and solubility in standard LC-MS solvent systems used for typical lipidomics measurements^58^. To resolve the issue of poor ionization, derivatization of sterols or the direct infusion of extracted lipidomes into ESI-MS instruments have been tried^59–61^, and while these methods have somewhat enabled the detection of cholesterol and its fatty acid esters, they require a dedicated standalone LC-MS instrument (cost extensive), are not amenable to other lipid measurements from the same biological samples, and in today’s age, are therefore not very popular with high-throughput lipidomics applications. To overcome the problem of solubility of sterols in conventional lipidomics solvents, numerous normal phase chromatographic techniques are reported using organic solvents such as hexanes, chloroform, tetrahydrofuran, isooctane^62–67^. Due the poor miscibility of these organic solvents with aqueous solvents, such normal phase lipidomics studies require special columns, LC-MS instruments, and are known to cause leaching of internal materials/tubing or clogging of seals for the LC system leading to increased system breakdowns over time. Despite resolving the issue of solubility of sterols, considering the problems associated with normal phase chromatography methods, and their incompatibility with conventional lipidomics strategies, both in terms of lipid extractions from biological samples and the downstream analysis, their use in high-throughput lipidomics applications remains limited.

Given the importance of cholesterol and its esters in mammalian biology, and the limited LC-MS lipidomics strategies reported to identify and quantify them from biological samples, we decided to develop a mass spectrometry-based method to bridge this gap. Here, we describe a simple, yet very sensitive LC-MS method that is compatible with conventional lipidomics techniques both in terms of lipid extractions from biological samples and the reverse-phase chromatography techniques (including solvent systems), which can robustly detect and quantify cholesterol and cholesteryl esters from various biological samples. Using this LC-MS method, we report the concentrations of cholesterol and its fatty acid esters in different mammalian cell lines and mouse tissues, and provide some insights into tissue-level cholesterol metabolism. Finally, we believe that the LC-MS method reported here, will provide the community with an invaluable, much-needed (conventional) lipidomics compatible mass spectrometry method for studying cholesterol metabolism in mammalian biology orthogonal to other cellular lipids under different physiological settings.

## MATERIALS AND METHODS

### Materials

Unless otherwise mentioned, all chemicals, buffers, and reagents were purchased from Sigma-Aldrich (Merck). All tissue culture media and supplies were purchased from HiMedia. All lipid standards used in this study were purchased from Avanti Polar Lipids. All solvents used for mass spectrometry were LC-MS grade and were purchased from JT Baker (methanol, MeOH) or Fisher Chemicals (isopropanol, IPA).

### Mammalian Cell Culture

All mammalian cell lines used in this study were purchased from ATCC and cultured in Dulbecco’s Modified Eagle’s Medium (DMEM) containing 10% (v/v) Fetal Bovine Serum (FBS) and 1% (v/v) Penicillin-Streptomycin (MP Biomedicals) at 37 °C and 5% (v/v) CO_2_. As a quality control step, all cell lines described in this study were routinely stained with 4’,6-diamidino-2-phenylindole (DAPI) and visualized by microscopy to ensure that they were devoid of any mycoplasma contamination^68, 69^. For cellular lipid extraction (per biological replicate), a single 10 cm tissue culture dish with cells at 80 – 85% confluence was used. Upon desired confluence, the cells were harvested by scraping, washed with cold sterile Dulbecco’s phosphate-buffered saline (DPBS) (three times), and finally centrifuged at 300g for 5 min at 4 °C to get a pellet. The harvested cell pellets were flash-frozen using liquid nitrogen and stored at −80 °C until further use.

### Tissue Samples from Mice

All protocols involving mice used for this study were approved by the Institutional Animal Ethics Committee at IISER Pune as per guidelines provided by the Committee for the Purpose of Control and Supervision of Experiments on Animals (CPCSEA) constituted by the Government of India (No: IISER_Pune IAEC/2023_03/02). All mice used in this study were from the C57Bl/6J strain and between 10 – 12 weeks of age. Briefly, the mice were deeply anesthetized, and whole blood samples (∼ 500 μL) were collected via the retro-orbital route into tubes containing 4% (w/v) ethylenediaminetetraacetic acid (EDTA) solution, immediately flash-frozen using liquid nitrogen and stored at −80 °C until further use. Following this, mice were sacrificed by cervical dislocation, and the various organs were harvested. All organs were washed with cold sterile DPBS (two times), transferred to 1.5 mL microcentrifuge tubes, weighed, flash-frozen using liquid nitrogen and stored at −80 °C until further use.

### Lipid Extraction from Mammalian Cells and Mouse Tissues

All lipid extractions were done using a modified Folch lipid extraction method, as per protocols previously reported by us^69–71^. Briefly, mammalian cell pellets were thawed on ice, re-suspended using 1 mL cold sterile DPBS, and transferred to a glass vial. On the other hand, the pre-weighed mouse tissues were transferred to Protein LoBind 1.5 mL microcentrifuge tubes (Eppendorf) using 500 µL cold sterile DPBS, and homogenized using Bullet Blender 24 (Next Advance) at an instrument speed setting of 8 for 3 min (two times). The resulting tissue homogenate was centrifuged at 500g for 5 min at 4 °C to remove tissue debris, and the supernatant was separated by gentle pipetting. The supernatant obtained was then made up to 1 mL with cold sterile DPBS and transferred into a glass vial. To the 1 mL of re-suspended cells or tissue homogenate (supernatant), 3 mL of 2:1 (v/v) chloroform:methanol (CHCl_3_: MeOH) containing the internal standard mix [1 nmol of C17:0 cholesteryl ester (catalog no.: 700186M); 2 nmol of cholesterol-d7 (catalog no.: 700041P)] was added. This mixture was vortexed vigorously and centrifuged at 1500g for 15 min to separate the mixture into two phases (bottom organic layer and top aqueous layer). The organic layer was removed by pipetting, transferred to a fresh glass vial, and dried under a stream of nitrogen gas. To remove any aqueous contaminants in the dried extract, it was resolubilized in 1 mL CHCl_3_, vortexed vigorously, transferred to a fresh glass vial, and dried again, under a stream of nitrogen gas. The final dried extracts were solubilized in 200 µL (for cells and whole blood) or 300 µL (for tissues) of 2:1 (v/v) CHCl_3_:MeOH, and 10 µL of this was used for the subsequent lipidomics (LC-MS) analysis.

### Liquid Chromatography-Mass Spectrometry (LC-MS)

Cholesterol and cholesteryl esters were quantified using the Auto-MS/MS acquisition mode on an Agilent 6545 Quadrupole Time Of Flight (QTOF) mass spectrometer fitted with an Agilent 1290 Infinity II UHPLC system. Reverse-phase liquid chromatography (LC) separation was achieved using a Gemini 5U C18 column (Phenomenex, 5 μm, 50 x 4.6 mm, catalog no.: 00B-4435-E0) coupled to a Gemini guard column (Phenomenex, 4 x 3 mm, catalog no.: AJ0-7597). The flow rate for the LC method was 0.5 mL/min, with the composition of Solvent A being 50:50 (v/v) H_2_O:MeOH + 0.1% formic acid + 10 mM ammonium formate, and that of Solvent B being 80:20 (v/v) IPA:MeOH + 0.1% (v/v) formic acid + 10 mM ammonium formate. A typical LC-run was 30 min, with the following gradients for elution of analytes of interest: 0 – 4 min, 40% solvent B; 4 – 6 min, 40 – 60% solvent B; 6 – 16 min, 60 – 100% solvent B; 16 – 22 min, 100% solvent B (wash step); 22 – 24 min, 100 – 40% solvent B; 24 – 30 min, 40% solvent B (re-equilibration step). The temperatures of the autosampler (samples) and column were maintained 10 °C and 40 °C respectively. All mass spectrometric acquisitions were performed using a dual Agilent jet stream electrospray ionization (AJS ESI) source in the positive ion mode, with the following MS parameters: drying and sheath gas temperature at 250 °C, drying and sheath gas flow at 10 L/min, nebulizer pressure at 45 psi, capillary voltage at 3 kV, nozzle voltage at 1 kV, and fragmentor voltage at 80 V. Precursor selections were carried out with a maximum of 4 precursor ions per cycle with collision energy set at 5 eV and using a preferred ion list optimized for cholesterol and cholesteryl esters.

### Data Analysis and Quantitation

To analyse the data from the LC-MS runs, using the METLIN Lipid library as a reference, a Personal Compound Database Library (PCDL) exclusively containing cholesterol and cholesteryl esters was curated using MassHunter PCDL manager B.08.00 (Agilent Technologies). The data files were processed in Agilent MassHunter Qualitative Analysis 10.0 using this PCDL library, and all the identified peaks were manually verified based on retention times of the LC-run and fragments obtained from the MS/MS spectral analysis. A mass accuracy threshold of 10 ppm was applied to all analytes for every experiment described in this study. The absolute concentrations of cholesterol or cholesteryl esters were quantified by measuring the area under the curve of an analyte relative to the respective internal standard and then normalizing this to the total protein content (cells or whole blood) or organ weight (tissues).

### Data Plotting

All data presented in this study are mean ± standard deviation from at least four biological replicates per experimental group. Unless otherwise mentioned, all graphs and plots reported in this study were made using the GraphPad Prism 10 [version 10.1.0 (264)] software for macOS.

## RESULTS

### Validation of LC-MS method

Our motivation in developing a conventional lipidomics compatible LC-MS method for detecting and quantifying cholesterol and cholesteryl esters stemmed from our interest in understanding the metabolism of cholesterol and its fatty acid esters in mammalian cells and tissues. For developing such a LC-MS method, we chose two commercially available unnatural sterol standards namely, cholesterol-d7 and cholesteryl heptadecanoate (C17:0 cholesteryl ester) (**Figure S1**) to standardize the LC elution profiles, and MS/MS fragmentation conditions. Both these sterol standards are near identical to endogenous cholesterol and cholesteryl esters, and can also be used as quantitative internal standards to estimate the concentrations of cholesterol and its fatty acid esters in lipid extracts prepared from different biological samples. Hence, they were chosen as starting candidates for optimizing the LC-MS method. It has been reported that commonly used ammonium additives in LC-MS solvents such as ammonium acetate or ammonium formate (important for charge generation on various neutral lipids), yield quasi-stable ammonium adducts [(M+NH_4_)^+^] for cholesterol in a positive ion mode LC-MS analysis. This ammonium adduct of cholesterol undergoes spontaneous dissociation (even at no collision energy in the MS) to generate a signature fragment ion of m/z 369.352 corresponding to a dehydrated form of cholesterol that is eventually detected in almost all LC-MS experiments done using ESI as a means for ionization (**Figure S2**)^58, 59, 64^. On the other hand, at zero collision energy conditions, the ammonium adducts for cholesteryl esters are known to be quite stable and are not prone to any form of dissociation (**Figure S2**)^58, 59, 64^.

With this knowledge, we injected a mixture of 1 nmol each of cholesterol-d7 and C17:0 cholesteryl ester and analysed the elution profile on column in our LC-MS method initially without the application of any collision energy in the MS. We specifically looked at the elution profiles for the mass to charge (m/z) values of 376.3956 and 656.6341 (with 10 ppm error), that corresponded to the dehydrated cholesterol-d7 species and the ammonium adduct of C17:0 cholesteryl ester respectively (**Figure 2A**). After performing this LC-MS analysis multiple times with the sterol standard mixture at 1 nmol of each, we always found single peaks for both m/z values, with the elution times of 11.7 ± 0.2 mins for m/z 376.3956 (with 10 ppm error) and 17.7 ± 0.2 mins for m/z 656.6341 (with 10 ppm error). Consistent with their respective hydrophobicity, we found that in our reverse-phased LC-MS method, the cholesterol standard eluted significantly before the C17:0 cholesteryl ester standard, and that overall, there was substantial separation in elution times of cholesterol versus the C17:0 cholesteryl ester in this LC-MS method.

Next, to determine the sensitivity and dynamic range of this LC-MS method, we performed a similar analysis using varying amounts of the two sterol standards. For cholesterol-d7, we performed this analysis from 10 pmol to 2 nmol range, while for C17:0 cholesteryl ester we used 1 pmol to 1 nmol range in this LC-MS method (**Figure 2B**). We found that within this range for cholesterol and cholesteryl esters, the LC-MS method has a linear dynamic range (over ∼ 3-orders of magnitude), and can reliably detect cholesterol and cholesteryl esters at 10 and 1 pmol respectively (**Figure 2B**), which is well below the known physiological concentrations for these lipids in mammalian cells and tissues.

**Figure 2.**
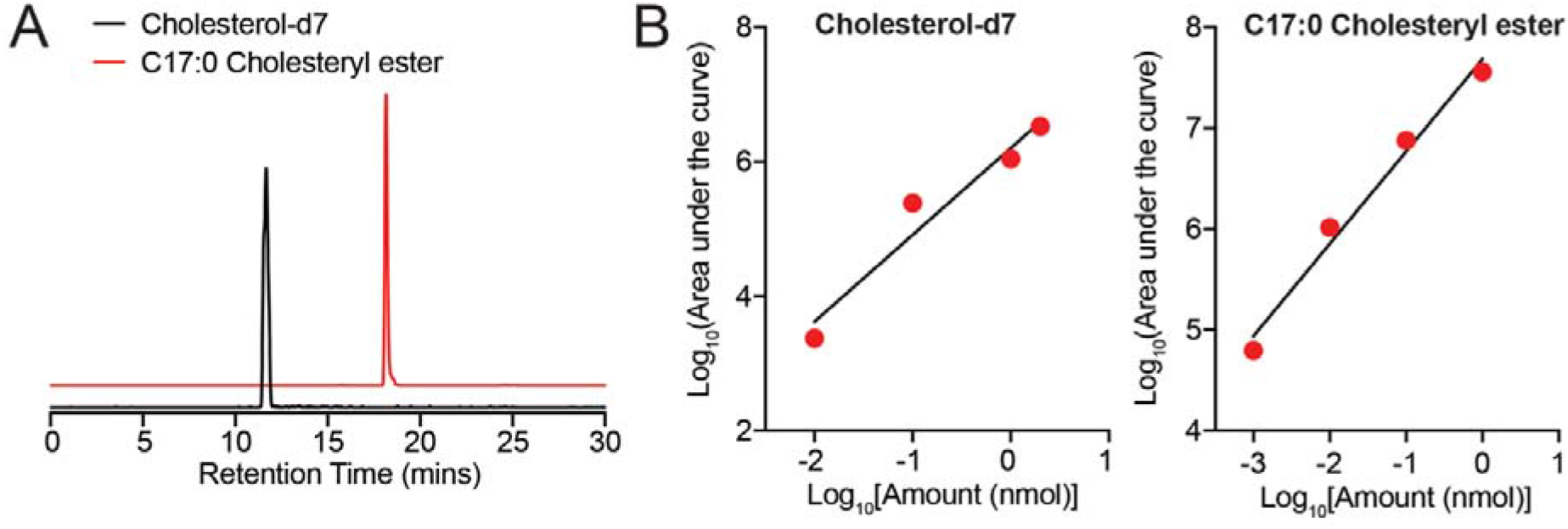
Elution and sensitivity of LC-MS method for detection of cholesterol-d7 and C17:0 cholesteryl ester. (**A**) Representative elution profiles for cholesterol-d7 (black trace, m/z = 376.3956 with 10 ppm error, 1 nmol) and C17:0 cholesteryl ester (red trace, m/z = 656.6341 with 10 ppm error, 1 nmol) in our LC-MS method. (**B**) Sensitivity and linear dynamic range for estimation of cholesterol-d7 and C17:0 cholesteryl ester in our LC-MS method.

Having established optimal elution conditions for cholesterol and cholesteryl esters in our LC-MS method, next, we wanted to obtain MS/MS fragments for these sterols, as this analysis would be essential and confirmatory, when assessing the levels of these lipids from complex biological samples. Toward this, we ran a mixture of 2 nmol cholesterol-d7 and 1 nmol C17:0 cholesteryl ester in our LC-MS method, now applying varying collision energies. We were looking for collision energy values where a significant of the parent ion would still be detected (needed for quantification of lipid to calculate area under the curve) and corresponding MS/MS fragments were also generated to confirm the identity of the sterol. From our analysis, we found that the collision energy of 5 eV was optimal for this, as below this, there was poor detection of the MS/MS fragments, and above this, the parent ion was almost fully converted to the MS/MS fragments. At a collision energy of 5 eV, cholesterol-d7 showed the parent ion (m/z = 376.396), and two dominant MS/MS fragments with m/z values of 283.242 and 256.219 (**Figure 3**), consistent with known fragments of cholesterol reported in literature^72^. Similarly, C17:0 cholesteryl ester showed the parent ion (m/z = 656.634) and a single MS/MS fragment of m/z = 369.351, that corresponds to the dehydrated form of cholesterol from the loss of the lipid tail upon application of 5 eV collision energy (**Figure 4**). Since, we could reliably identify signature MS/MS fragments for both cholesterol and cholesteryl esters, we chose 5 eV as the collision energy in our LC-MS method, and used these conditions to quantitatively profile cholesterol and its different fatty acid esters in mammalian cells and tissues.

**Figure 3.**
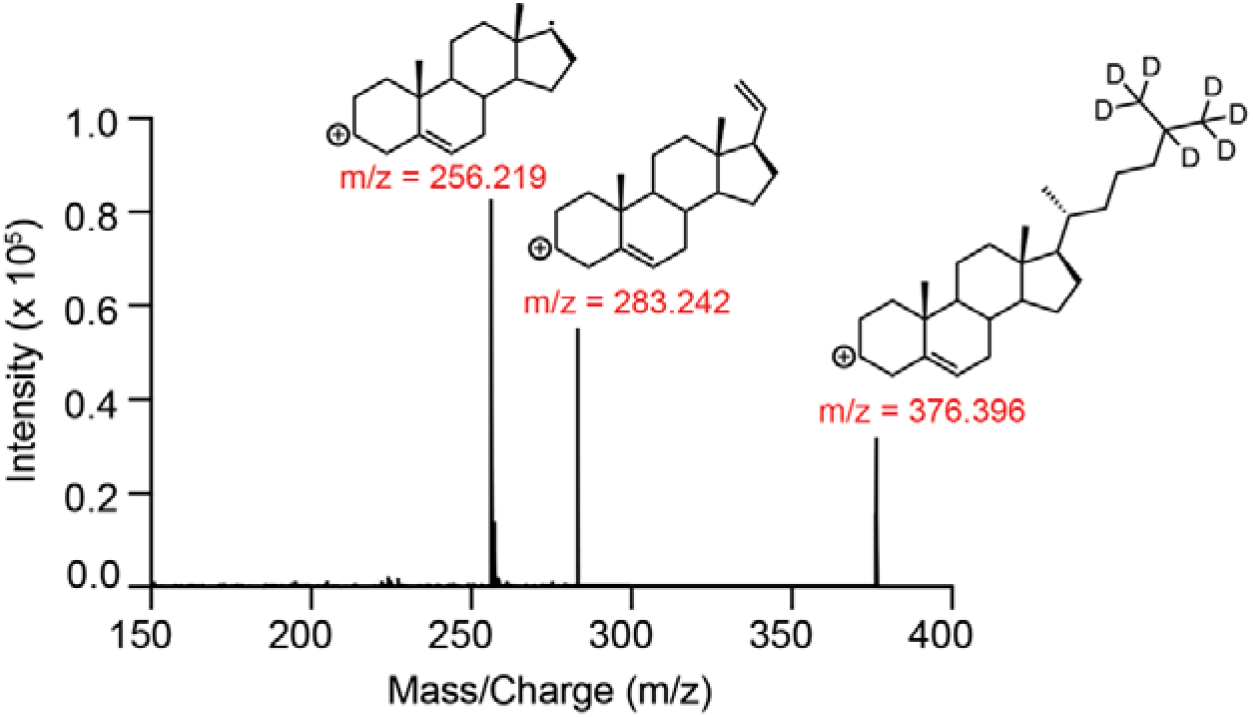
MS/MS analysis of cholesterol-d7. 2 nmol of cholesterol-d7 was run in our LC-MS method with a collision energy of 5 eV. Besides the parent ion (m/z = 376.396 for cholesterol-d7), two signature MS/MS fragments can be used to confirm the presence of cholesterol.

**Figure 4.**
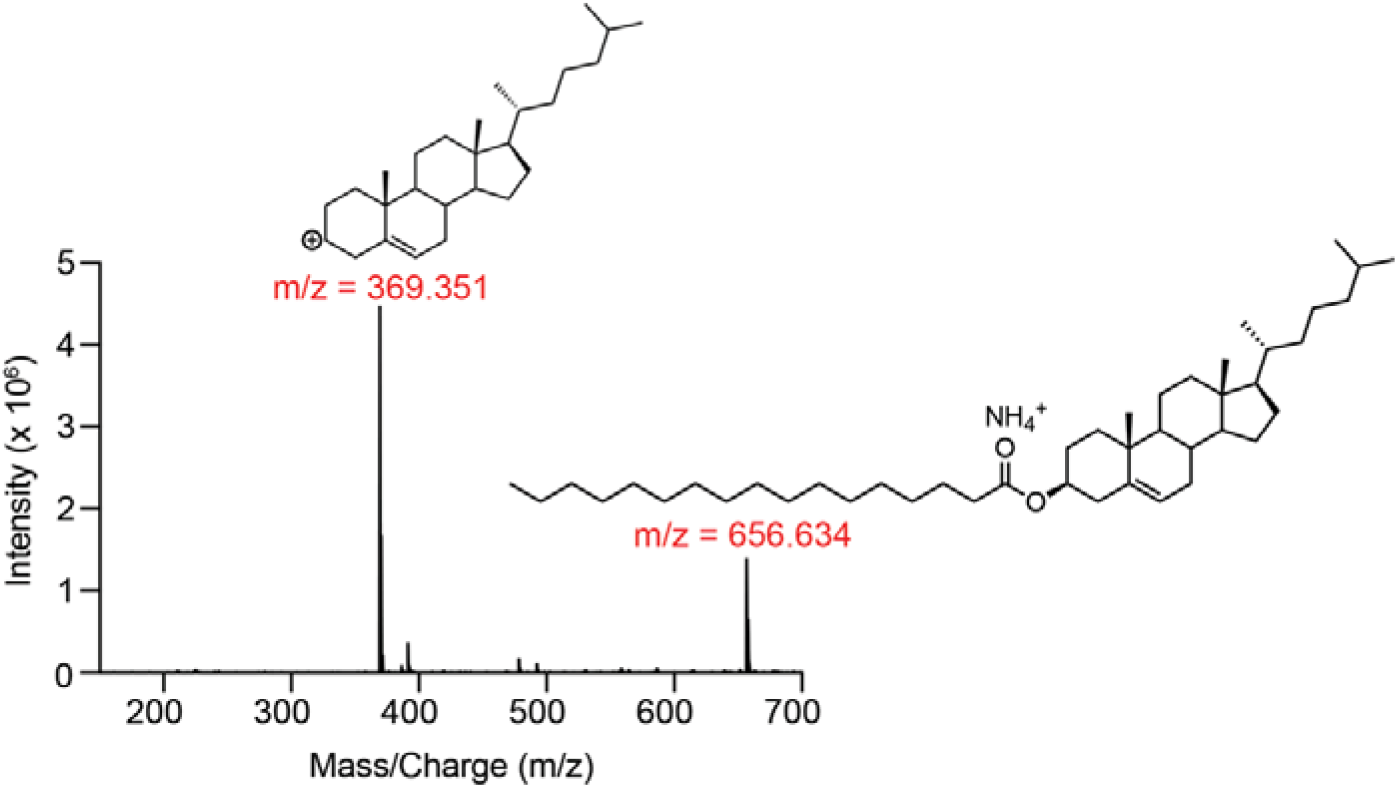
MS/MS analysis of C17:0 cholesteryl ester. 1 nmol of C17:0 cholesteryl ester was run in our LC-MS method with a collision energy of 5 eV. Besides the parent ion (m/z = 656.634 for C17:0 cholesteryl ester), a single signature MS/MS fragment (m/z = 369.351) corresponding to dehydrated cholesterol can be used to confirm the presence of cholesteryl esters.

### Measurement of cholesterol and cholesteryl esters in mammalian cells

Having developed a LC-MS method to study cholesterol and cholesteryl esters, next we wanted to assess the capability of this method to reliably detect and quantify these lipids in different mammalian cell lines. For this, we chose three immortalized cell lines, namely HEK293T (general mammalian cells), RAW264.7 (macrophage cell line) and Neuro2A (neuronal cell line), as cholesterol metabolism is known to be important in the immune and nervous system^1, 2, 24, 26, 33^. Since the eventual goal of this method was the need to be compatible with existing high-throughput lipidomics LC-MS platforms, we decided to use the same lipid extraction protocols (reported by others and us^69–71^) that have already been set in place for such experiments. Based on this lipidomic extraction, we set up a workflow for measuring cholesterol and its fatty acid esters in the aforementioned mammalian cells (and also tissues, discussed in next section) (**Figure 5**).

**Figure 5.**
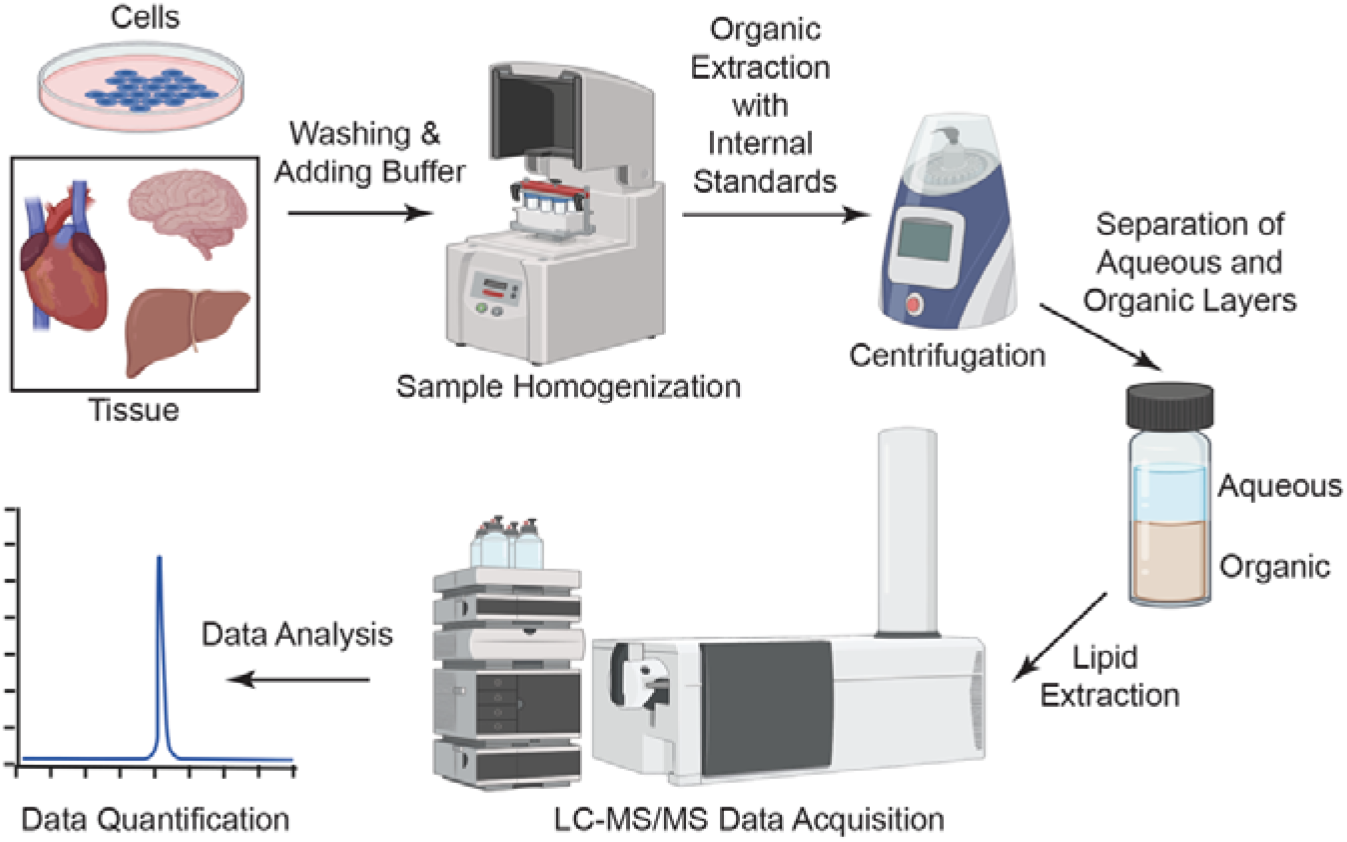
A general workflow of extracting lipids and measuring cholesterol and its esters from mammalian cells and tissues.

Following this workflow, our analysis showed that across these three mammalian cell lines, we could robustly and reproducibly detect both cholesterol and its fatty acid esters. Specifically, we found that relative to HEK293T or Neuro2A cells, RAW264.7 cells have significantly more cholesterol (**Figure 6**). This is expected, as monocytic cells (like macrophages) generally tend to accumulate cholesterol, which they require to orchestrate immune responses, and our cholesterol measurements data supports this notion^33, 73, 74^. Next, we analysed the cholesteryl ester levels from these three mammalian cell lines and found that the different cell lines have distinct distributions of cholesteryl esters (**Figure 6, Table S1**). Interestingly, we found that compared to saturated variants, all these cell lines had significantly higher concentrations of mono-(e.g. cholesteryl oleate, C18:1) or polyunsaturated (e.g. cholesteryl linoleate, C18:2) variants of cholesteryl esters. Our measurements also showed that certain cell lines had a preference for specific cholesteryl esters, for example, RAW264.7 preferred cholesteryl oleate, while Neuro2A cells preferred cholesteryl linoleate. While the physiological significance of why certain cell lines prefer certain variants of cholesteryl esters remains unknown, our LC-MS method shows for the first time, the complete diversity of cholesteryl esters present across different mammalian cell lines, as most studies to date only report the abundant cholesteryl esters.

**Figure 6.**
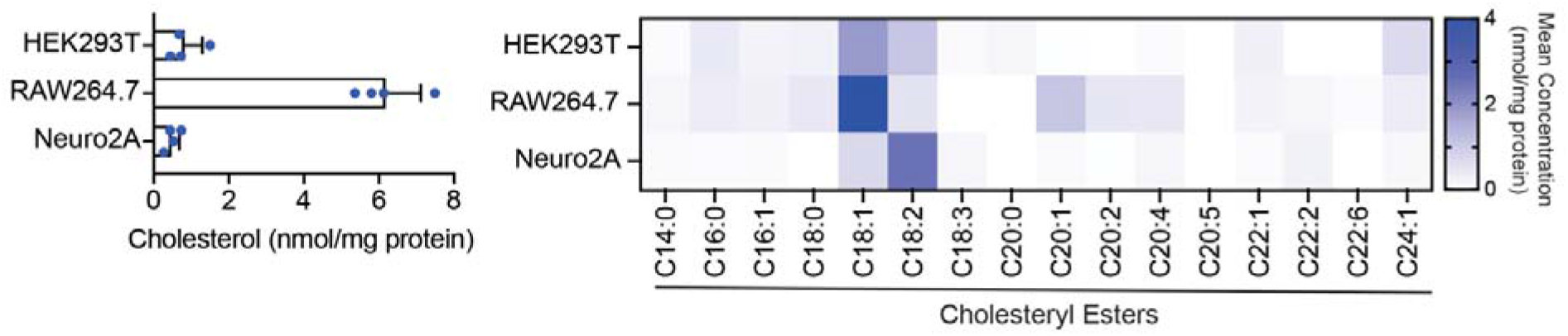
Measurements for cholesterol and cholesteryl esters in different mammalian cell lines. *<u>Left panel</u>* shows the cholesterol concentrations in different mammalian cell lines. Bar data is represented as mean ± standard deviation from four biological replicates. *<u>Right panel</u>* shows the concentrations of diverse cholesteryl esters in the different mammalian cell lines. Heat map data is represented as mean of concentration from four biological replicates. Individual data for different cholesteryl esters in the cell lines can be found in **Table S1**.

### Measurement of cholesterol and cholesteryl esters in mouse tissues and blood

Having profiled cholesterol and its fatty acid esters in different mammalian cell lines, we next attempted to measure these lipids in various mouse tissues using the same LC-MS method. For this experiment, we harvested different mouse tissues and blood, extracted lipids from them using established high-throughput lipidomics compatible protocols^69–71^, and analysed the cholesterol and cholesteryl ester content of these lipid extracts using our LC-MS method. Surprisingly, we found that different tissues had very distinct cholesterol and cholesteryl ester profiles, and our data might provide some interesting insights in the metabolism of these lipids in mammals (**Figure 7, Table S1**). For instance, we found that the brain had the highest cholesterol concentration, and perhaps the lowest concentrations of cholesteryl esters amongst all the tissues. This data fits well with reports which suggest that given the need for cholesterol (not its fatty acid esters) during neuronal myelination, there is substantial *de novo* cholesterol biosynthesis in the mammalian brain^31, 32, 75^. On the contrary, in analysis of the blood from mice, consistent with heightened blood LCAT activity^18–20^, we found that systemically circulating cholesterol is sequestered, converted to HDLs, and stored in various tissues for later use.

**Figure 7.**
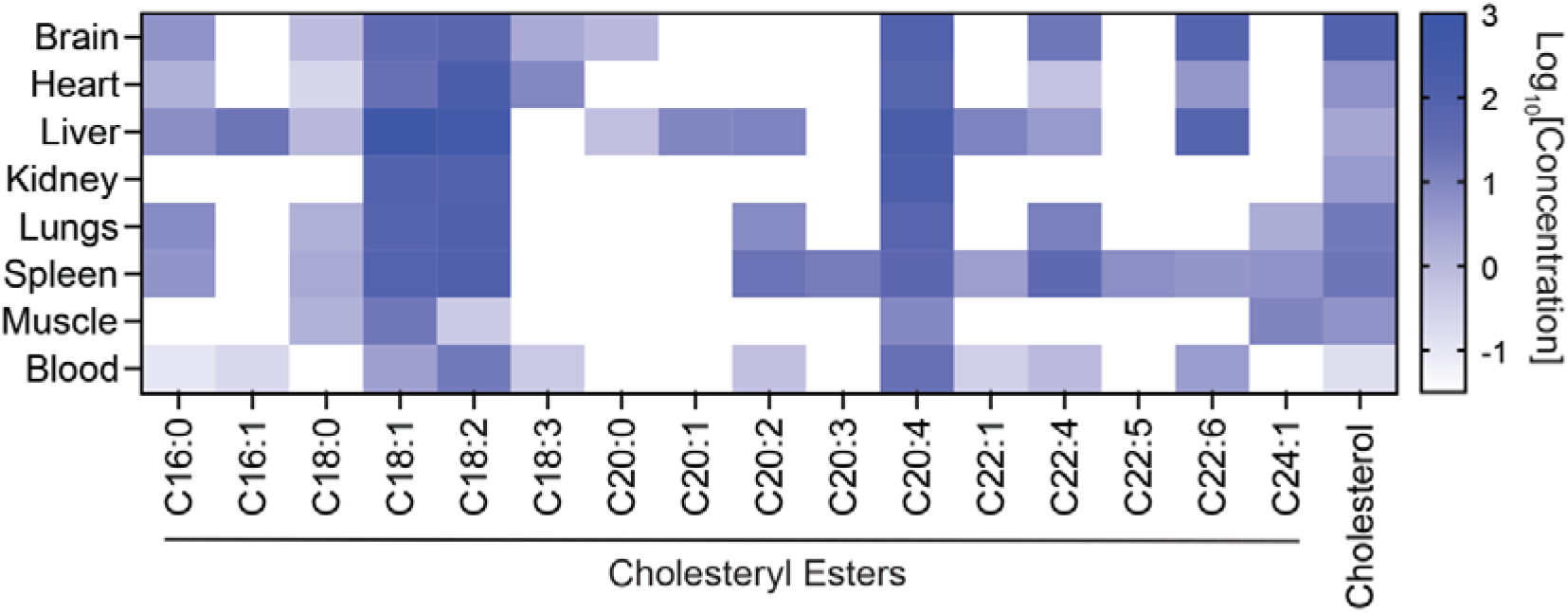
Concentration of cholesterol and cholesteryl esters in different mouse tissues and blood. Heat map data is represented in logarithmic scale (to base 10) of the mean concentration from four biological replicates. The units for concentration of cholesterol and cholesteryl esters in tissues are nmol/g tissue weight, while that in blood is nmol/mg protein. Individual data for cholesterol and the different cholesteryl esters in mouse tissues can be found in **Table S1**.

More generally, we found that most tissues (except brain and blood) had comparable concentrations of cholesterol (**Figure 7**, **Table S1**). Amongst cholesteryl esters, we found that monounsaturated (e.g. cholesteryl oleate (C18:1)), and polyunsaturated [e.g. cholesteryl linoleate (C18:2) and cholesteryl arachidonate (C20:4)] variants were the most abundant species, while the saturated variants [e.g. cholesteryl palmitate (C16:0) and cholesteryl stearate (C18:0)] had much lower concentrations across all tissues (**Figure 7, Table S1**). Not surprisingly, we found that the liver had the highest tissue concentration and diversity of cholesteryl ester variants amongst the different tissues profiled, which is consistent with its known roles in cholesterol transport and metabolism^2, 24, 26, 76, 77^. Interestingly, we found that despite its active role in regulation of cholesterol flux, the kidney had the least diversity of cholesteryl esters, and possessed only the aforementioned monounsaturated and polyunsaturated cholesteryl ester variants (**Figure 7, Table S1**). Taken together, after profiling different mammalian tissues, using extracts from standard lipidomics extractions, the LC-MS method developed here, can quantitatively profile cholesterol and 19 fatty acid esters of cholesterol in tandem (**Table 1**). This study for the first time, provides a comprehensive picture of cholesterol and cholesteryl esters in different mouse tissues under *ad libitum* feeding (normal physiological) conditions.

**Table 1:** Information of different cholesterol and cholesteryl esters detected in mammalian cells and tissues.

| Species | Parent Ion<br>m/z | Product Ion<br>m/z | Species<br>Detected | Retention Time<br>(min) |
| --- | --- | --- | --- | --- |
| Cholesterol | 369.3516 | 256.2186 | $[M-H_2O+H]^+$ | $11.7 \pm 0.2$ |
| **Cholesterol-d7**<br>**(Internal standard)** | 376.3956 | 256.2186 | $[M-H_2O+H]^+$ | $11.7 \pm 0.2$ |
| C14:0 cholesteryl ester | 614.5872 | 369.3516 | $[M+NH_4]^+$ | $15.6 \pm 0.1$ |
| C16:0 cholesteryl ester | 642.6185 | 369.3516 | $[M+NH_4]^+$ | $16.8 \pm 0.1$ |
| C16:1 cholesteryl ester | 640.6029 | 369.3516 | $[M+NH_4]^+$ | $16.5 \pm 0.1$ |
| C16:2 cholesteryl ester | 638.5872 | 369.3516 | $[M+NH_4]^+$ | $16.2 \pm 0.1$ |
| **C17:0 cholesteryl ester**<br>**(Internal standard)** | 656.6342 | 369.3516 | $[M+NH_4]^+$ | $17.7 \pm 0.2$ |
| C18:0 cholesteryl ester | 670.6498 | 369.3516 | $[M+NH_4]^+$ | $18.0 \pm 0.2$ |
| C18:1 cholesteryl ester | 668.6342 | 369.3516 | $[M+NH_4]^+$ | $17.6 \pm 0.2$ |
| C18:2 cholesteryl ester | 666.6185 | 369.3516 | $[M+NH_4]^+$ | $17.0 \pm 0.2$ |
| C18:3 cholesteryl ester | 664.6029 | 369.3516 | $[M+NH_4]^+$ | $16.6 \pm 0.2$ |
| C20:0 cholesteryl ester | 698.6811 | 369.3516 | $[M+NH_4]^+$ | $18.2 \pm 0.2$ |
| C20:1 cholesteryl ester | 696.6655 | 369.3516 | $[M+NH_4]^+$ | $18.0 \pm 0.2$ |
| C20:2 cholesteryl ester | 694.6498 | 369.3516 | $[M+NH_4]^+$ | $17.8 \pm 0.2$ |
| C20:4 cholesteryl ester | 690.6185 | 369.3516 | $[M+NH_4]^+$ | $17.6 \pm 0.2$ |
| C20:5 cholesteryl ester | 688.6029 | 369.3516 | $[M+NH_4]^+$ | $17.3 \pm 0.2$ |
| C22:1 cholesteryl ester | 724.6968 | 369.3516 | $[M+NH_4]^+$ | $18.9 \pm 0.2$ |
| C22:2 cholesteryl ester | 722.6811 | 369.3516 | $[M+NH_4]^+$ | $18.6 \pm 0.2$ |
| C22:4 cholesteryl ester | 718.6498 | 369.3516 | $[M+NH_4]^+$ | $18.3 \pm 0.2$ |
| C22:5 cholesteryl ester | 716.6342 | 369.3516 | $[M+NH_4]^+$ | $17.9 \pm 0.2$ |
| C22:6 cholesteryl ester | 714.6185 | 369.3516 | $[M+NH_4]^+$ | $17.7 \pm 0.2$ |
| C24:1 cholesteryl ester | 752.7281 | 369.3516 | $[M+NH_4]^+$ | $19.2 \pm 0.2$ |

## DISCUSSION

In mammals, cholesterol is an indispensable neutral lipid that serves many critical physiological functions, and often serves as a lynchpin in sterol/fat metabolism^1, 2, 24, 26^. During this metabolism, it is known that excess free cholesterol is stored as comparatively inert cholesteryl esters, and these fatty acid esters of cholesterol are known to be present in almost all organs in mammals (**Figure 1**). Given their importance in mammalian physiology and strong associations with human diseases^25–34^, over the years, numerous attempts have been made to develop analytical methods to detect and quantify cholesterol and its fatty acid esters in biological samples. In fact, a few of these analytical methods are currently used in pathological labs worldwide to report cholesterol (but not cholesteryl esters) and lipoprotein levels from human blood samples^35^. Yet, despite this progress, there are virtually no reliable LC-MS methods available that are compatible with currently used high-throughput lipidomics methodologies, which can profile cholesterol and its fatty acid esters in tandem with other extracted lipids from biological samples using reverse-phase chromatographic techniques. This is particularly important because pathophysiological conditions like dyslipidemia or obesity are complex metabolic states, where multiple lipid pathways are dysregulated in tandem. However, given the technical limitations mentioned earlier, it has been a challenge to understand how sterol metabolism crosstalks with or affects other lipid pathways during such metabolic conditions from current high throughput LC-MS based lipidomics experiments.

Realizing this, we attempted to develop a LC-MS method that was compatible with existing lipidomics strategies and, capable of identifying and quantifying cholesterol and cholesteryl esters from mammalian cells and tissues. Here, we describe a reverse-phase LC-MS method which is both sensitive and robust in quantifying cholesterol and its fatty acid esters using specific MS/MS fragments of these lipids from extracts obtained using standard organic extraction protocols^69–71^ (**Figures 2–5**). Using this LC-MS method, we quantify cholesterol and cholesteryl esters in mammalian cells (**Figure 6**) and mouse tissues and blood (**Figure 7**). Our studies show that cholesterol is ubiquitous in all mammalian cell lines and tissues (including blood), but its concentrations vary significantly across these biological samples. We find that amongst cholesteryl esters, the monounsaturated (specifically C18:1), and polyunsaturated (specifically C18:2 and C20:4 (in tissues only)) variants were the most abundant in different mammalian cells and tissues. We also find that saturated variants have fairly low abundance in mammalian cells and tissues, while on the other hand, we are able to robustly detect an array of very-long chain (≥ C20) polyunsaturated cholesteryl esters that have proved challenging to measure in the past, with differing concentrations in different mammalian tissues (**Table 1, Table S1**).

The LC-MS method developed by us has several advantages. First, and most important, this method does not require the chemical derivatization of cholesterol or cholesteryl esters, and can robustly detect these lipids in lipid extracts of mammalian cells and tissues obtained from standard organic extraction protocols^69–71^. Therefore, this method can reliably measure both cholesterol and its fatty acid esters in tandem from the very same lipidomic preparations used to profile other lipids (e.g. phospholipids, sphingolipids, neutral glycerides etc.) and provide direct information on cholesterol metabolism relative to these lipids. Second, this method is robust, sensitive, and can reliably quantify the physiological concentrations of both cholesterol and cholesteryl esters from diverse biological samples. Third, here, we describe this LC-MS method using a QTOF mass spectrometer. However, given the information provided by us in terms of LC-solvent system or column, and the MS/MS fragments provided for cholesterol and its fatty acid esters, we are confident that this reverse-phase LC-MS method can easily be adapted to other ESI-MS systems (e.g. Triple Quadrupole, QTRAP, Orbitrap etc.) for quantifying cholesterol and cholesteryl esters.

Moving ahead, this LC-MS method can be used to study several aspects of cholesterol metabolism. For example, while significant knowledge is available on the conversion of cholesterol to cholesteryl esters by ACATs and LCAT, the identity of the enzymes that degrade cholesteryl esters back to cholesterol in mammalian cells and tissues still remain cryptic and somewhat controversial^22, 23, 78–81^. Our data suggests that most tissues have substantial amounts of stored cholesterol in the form of cholesteryl esters, particularly the monounsaturated and polyunsaturated variants, and therefore, there must exist lipases capable of metabolizing these fatty acid esters of cholesterol to maintain homeostatic levels of free cholesterol under varying physiological conditions. To this end, our LC-MS method can be used as a reliable readout of these lipids, and complement *in vitro* pharmacological screens, or *in vivo* genetic or chemical genetic screens to identify such lipases (cholesterylesterases) in different mammalian cells and tissues. Further, it is known that under conditions of oxidative stress, there are enzymatic and non-enzymatic pathways in humans that convert cholesterol and its fatty acid esters to potent immunogenic lipid mediators such as oxysterols^82–85^ and oxidized cholesteryl esters^86–89^. Given their emerging roles in mammalian immunology, over the years, there have been some efforts at looking at these oxidatively modified variants of cholesterol and its fatty acid esters using various LC-MS techniques^62, 63, 90–93^. Yet, most of these LC-MS methods are not compatible with existing high-throughput lipidomics strategies. We believe that the LC-MS method developed by us, can also be modified to detect and quantify such oxidatively modified variants of cholesterol and cholesteryl esters from mammalian cells and tissues under physiological conditions that involve oxidative stress.

## CONCLUSION

Over the years quantifying cholesterol and cholesteryl esters from biological samples using LC-MS analysis has proved rather difficult owing to bottlenecks from sample preparations, efficient separation and ionization of these lipids. Here, we developed a much-needed robust high-throughput lipidomics compatible LC-MS method for efficiently identifying and quantifying cholesterol and cholesteryl esters from biological samples. Using this method, we successfully profiled cholesterol and its fatty acid esters in a few mammalian cell lines and different mouse tissues and blood. Our studies provide the first comprehensive picture of the diversity amongst this lipid class in mammals under physiological conditions. We believe that the LC-MS method reported here will greatly facilitate the study of cholesterol and its fatty acid esters in mammalian biology, and open new avenues for understanding their functional roles in as-of-yet poorly characterized pathophysiological conditions and human diseases.

## AUTHOR CONTRIBUTIONS

A.C. performed all the experiments. A.C. and S.S.K. conceived the project, analyzed all the data and wrote the paper. S.S.K. supervised and acquired funding for the project.

## CONFLICT OF INTEREST

The authors declare no competing financial interests.

## FUNDING

This study was supported by funds from a SwarnaJayanti Fellowship (to S.S.K.) from the Science and Engineering Research Board (SERB), Department of Science and Technology, Government of India (grant number: SB/SJF/2021-22/01). A.C. is supported by a graduate student fellowship from IISER Pune. The mice used in this study were maintained and provided by the National Facility for Gene Function in Health and Disease (NFGFHD) at IISER Pune, which was supported by a grant from the Department of Biotechnology, Govt. of India (grant number: BT/INF/22/SP17358/2016).

## ACKNOWLEDGEMENT

Members of the S.S.K. lab are thanked for providing critical comments and discussions throughout the course of this study. We thank Saddam Shekh for technical assistance and maintenance of the LC-MS facility at IISER Pune, and the staff members of the NFGFHD at IISER Pune for helping with the mice experiments described in this study.

## Notes

### Competing Interest Statement

The authors have declared no competing interest.

