## Supplementary Figures for "A Facile LC-MS Method for Profiling Cholesterol and Cholesteryl Esters in Mammalian Cells and Tissues"

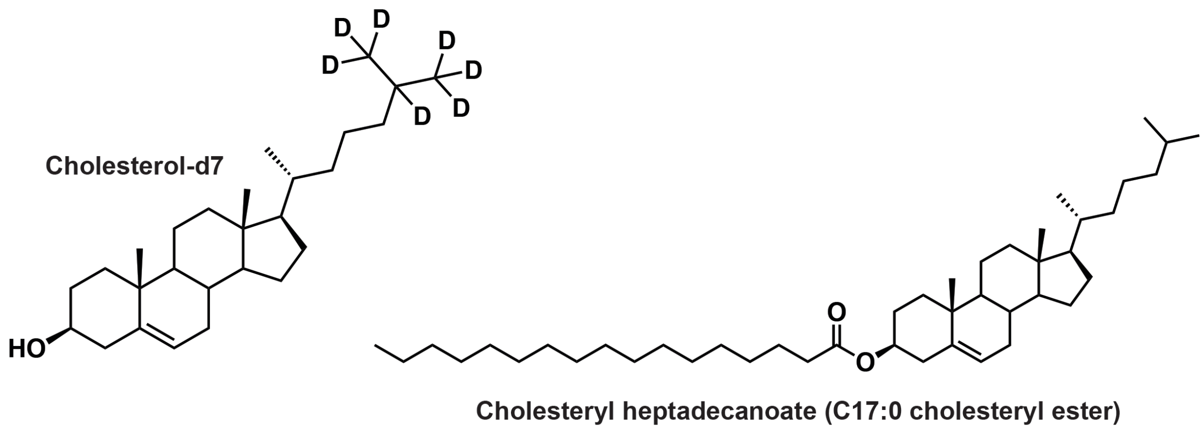


**Figure S1.** Structures of cholesterol-d7 and C17:0 cholesteryl ester that are used as internal standards for quantification of cholesterol and cholesteryl esters respectively in our LC-MS method.

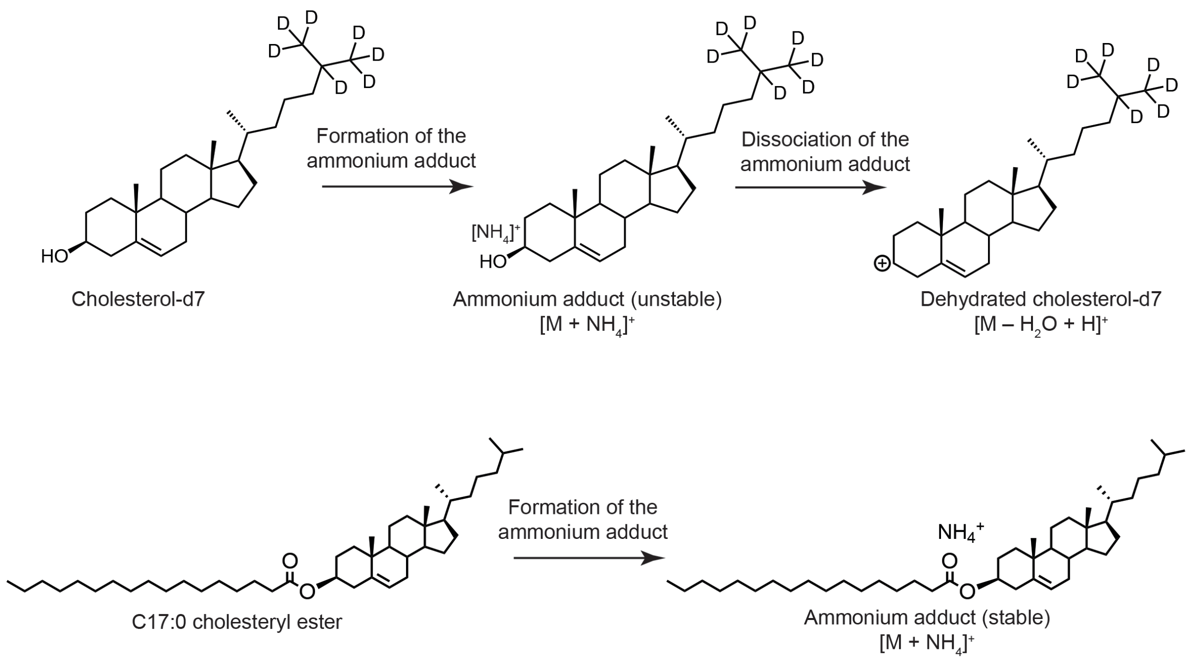


**Figure S2.** Formation of ammonium adducts of cholesterol-d7 and C17:0 cholesteryl ester in the ESI-MS instrument, and the eventual parent ions detected.
